# Loss of the first β-strand of human prion protein generates an aggregation-competent partially “open” form

**DOI:** 10.1101/2022.09.20.508729

**Authors:** Laszlo L. P. Hosszu, Daljit Sangar, Mark Batchelor, Emmanuel Risse, Andrea M. Hounslow, Jonathan P. Waltho, John Collinge, Jan Bieschke

## Abstract

Prion diseases, a group of incurable, lethal neurodegenerative disorders of mammals including humans, are caused by prions, assemblies of misfolded host prion protein (PrP). The pathway of PrP misfolding is still unclear, though previous data indicate the presence of a structural core in cellular PrP (PrP^C^), whose cooperative unfolding presents a substantial energy barrier on the path to prion formation. PrP is a GPI-anchored membrane protein, and a number of studies suggest that membrane interactions play an important role in the conversion of PrP^C^ to its disease-associated form, including a transmembrane form of PrP in which a highly conserved region (residues 110 - 136) spans the ER membrane. Insertion of this region results in the detachment of the PrP^C^ first β-strand from the structural core. The effect of this removal on the structure, stability and self-association of the folded domain of PrP^C^ is determined here through a biophysical characterisation of a truncated form of PrP^C^ lacking this region. Whilst markedly destabilised, NMR chemical shifts show that the truncated protein exhibits tertiary structure characteristic of a fully folded protein and retains its native secondary structure elements, including the second strand of the PrP β-sheet, but with altered conformational flexibility in the β2-α2 loop and first α-helix. The latter is destabilised relative to the other helical regions of the protein, with markedly increased solvent exposure. This truncated form of PrP fibrilises more readily than the native form of the protein. These data suggest a stepwise mechanism, in which a destabilised “open” form of PrP^C^ may be a key intermediate in the refolding to the fibrillar, pathogenic form of the protein.

## Introduction

Prion diseases, such as bovine spongiform encephalopathy (BSE) in cattle, scrapie in sheep, and Creutzfeldt-Jakob disease (CJD), kuru, and Gerstmann-Sträussler-Scheinker (GSS) syndrome in humans, are a group of neurodegenerative disorders caused by prions, self-replicating β-sheet-rich infectious polymeric assemblies of misfolded host-encoded cellular prion protein (PrP^C^) (1-6). Whilst rare, prion diseases are an area of intense research interest, as it is increasingly recognised that other degenerative brain diseases, such as Alzheimer’s and Parkinson’s diseases, also involve the accumulation and spread of aggregates of misfolded host proteins through an analogous process of seeded protein polymerisation (2, 7-10). Consequently, study of ‘prion-like’ mechanisms has been recognised to have much a wider relevance to the understanding of neurodegenerative disorders (11-13).

PrP^C^ is a widely-expressed cell surface, glycosylphosphatidylinositol (GPI)-anchored glycoprotein that is sensitive to protease treatment and is soluble in detergents. PrP may have a role in cell adhesion or signalling processes, but its precise cellular function remains unknown. It consists of two structural domains, an unstructured N-terminus, spanning approximately residues 23 - 125, which contains five repeats consisting of a single nonapeptide and four octapeptides, and a structured, mainly α-helical C-terminal domain, which includes a single disulphide bond and two glycosylation sites (1, 14, 15). A highly conserved section, (approximately residues 110-136) referred to as the conserved hydrophobic region (CHR) spans both domains. Prions, in contrast, are dominated by β-sheet structure and are insoluble in detergents, with some displaying marked protease-resistance. These have been classically designated as PrP “Scrapie” (PrP^Sc^) and are found only in prion-infected tissue (16).

Prion diseases may be acquired (transmitted between animals or humans), inherited or sporadic (of unknown cause). Inherited prion diseases comprise around 10-15% of total prion disease cases, with over 30 different pathogenic somatic mutations and approximately 10 polymorphic variants having been identified in the human PrP gene (*PRNP*) (17). Amyloid formation in a range of proteins is associated with destabilisation of the protein native state through somatic mutations (18-21). Although destabilisation of PrP^C^ does not correlate with specific disease phenotypes and no definitive link between PrP stability and disease has been established, the majority of *PRNP* pathogenic mutations reside within the structured C-terminal domain, with all fully penetrant pathogenic mutations showing significant destabilisation (22). *In vitro* experiments also show that destabilisation of native PrP^C^ through the use of chemical denaturants, temperature, redox conditions, pH or nucleic acid-binding favours the formation of protease-resistant amyloid or fibrillar structures, some of which have been claimed to be associated with disease (23-25). The most general model proposed thus far for the process of nucleated protein polymerisation is that PrP^C^ fluctuates between its dominant native state and other minor isoforms. These can self-associate in an ordered manner to produce a stable supramolecular structure composed of misfolded PrP monomers, which can convert other isoforms to the infectious isomer in an autocatalytic manner (2).

Many studies have thus focused on characterising partially folded intermediate states which may be involved in PrP self-association. A key observation in this regard is that the three α-helices and the second strand of the PrP^C^ β-sheet form a stable assembly, which may be considered to be the core of the protein. This is because these structural elements have stabilities equivalent to the overall fold stability of the protein and the backbone amides of these structural elements exchange with solvent solely when PrP^C^ completely unfolds (26). Minor isoforms which retain this structural core are thus energetically favoured over those in which this core region is disrupted. Notably, the first strand of the PrP β-sheet, which is also the most N-terminal structural element in the folded C-terminal domain, lies outside of this core region and displays markedly reduced stability in comparison to the other PrP secondary structure elements (approximately 30-fold less) (26). This raises the possibility that this secondary structure element may act independently of the core of PrP^C^, and indeed NMR chemical shift data indicate that the PrP^C^ native state ensemble contains a population of conformers where the PrP N-terminus, including the first strand of the PrP^C^ β-sheet, is detached from the PrP^C^ core (27).

The detached region comprises up to residue 146, and includes the conserved hydrophobic region of PrP (CHR), which spans residues 110 – 136. The CHR displays exceptionally high conservation across a wide range of species, indicative of an essential role in the endogenous function of PrP^C^ (28, 29). The CHR controls PrP co-translational translocation at the endoplasmic reticulum (ER) during the biosynthesis of the protein (30). It can associate with the ER membrane in various topologies, with mutations in this region upregulating particular PrP isoforms, which can lead to neurodegenerative disease in mice and some heritable prion diseases (31). Its amino acid sequence also displays many of the characteristics of a transmembrane helix (32), and NMR studies show that residues within this region interact with membrane analogues (dodecylphosphocholine micelles) in aqueous solution both as part of PrP^C^ (33) and as a distinct peptide (29). Here, it adopts a membrane-spanning helical conformation when associated with lipid micelles (29) and a radically different conformation to the reported solution structures of PrP^C^. In addition, antibody binding to the CHR, which would inhibit membrane insertion, also inhibits the propagation of proteinase K-resistant PrP^Sc^ and prion infectivity, further implicating the CHR and its membrane association in prion disease (34). Competitive antibody binding to the CHR has been proposed to block interactions between PrP and cofactor molecules thereby inhibiting PrP conversion and prion propagation (34).

Insertion of the CHR into the endoplasmic reticulum (ER) membrane necessarily involves detachment of residues 110 - 136 from the folded C-terminal domain, including the first strand of the PrP β-sheet, from the PrP^C^ core. Whilst the proportion of PrP molecules with a detached CHR region is low in solution (27), a high local concentration of membrane-binding sites for membrane-incorporated PrP would be available, likely resulting in a significant bound population. Thus, it is reasonable to consider whether these membrane-associated forms of PrP^C^ may destabilise the PrP^C^ core region and facilitate its self-association. Our aim in this study was to investigate the effect of the removal of the CHR on the structure, stability and self-association of the remaining folded domain of PrP, through the biophysical characterisation of truncated PrP^C^ lacking the CHR. We find that this form of the protein whilst retaining many of the characteristics and structure elements of native PrP^C^, has substantially increased solvent exposure relative to the native state, and is significantly more susceptible to fibrillisation, raising the possibility that this partially-folded state may be a principal precursor in the formation of ordered fibrillar structures.

## Results

### Choice of PrP constructs studied

In order to assess the effects of removal of the CHR on the structured domain of PrP^C^ we generated a recombinant PrP construct spanning residues 137-231, which we termed PrP^137^. Data for this construct was primarily compared with that of PrP^119^ (residues 119-231 of PrP), which contains the structured elements of PrP^C^, and does not contain an extended unstructured N-terminus, which complicates the structural analyses.

### PrP^137^ retains native-like secondary and tertiary structure

Circular dichroism (CD) and NMR spectra of PrP^137^ in solution were acquired to provide information on the secondary and tertiary structure of the protein. The near-UV CD spectrum of PrP^137^ (Fig. 2 (a)) is broadly similar to that of PrP^119^, with two minima at approximately 208 and 222 nm, characteristic of a predominantly α-helical protein. The loss of β-sheet signal resulting from the loss of residues 119-136 is masked by the significantly more intense α-helical signal. PrP^137^ thus appears to retain the mainly α-helical structure of the untruncated C-terminal domain. There is some reduction in the mean residue ellipticity (MRE) for the minimum at 208 nm, which is suggestive of some alteration in the packing of the α-helices however.

**Figure 1.**
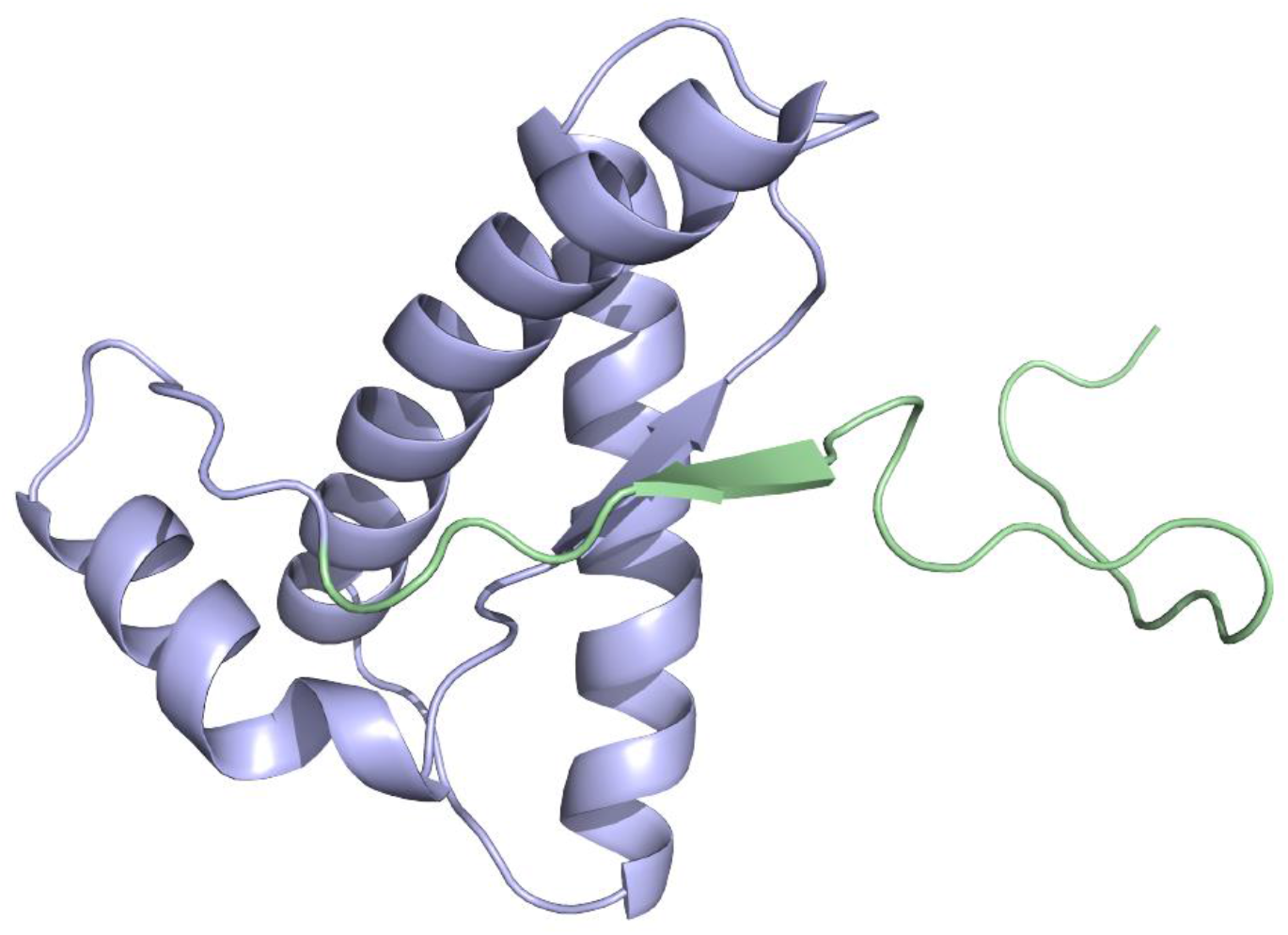
Cartoon representation of PrP^C^ secondary structure (14, 15). Residues 119-226 are displayed, with the N-terminal region deleted in PrP^137^ highlighted in green. The first strand of the PrP β-sheet is removed in PrP^137^.

**Figure 2.**
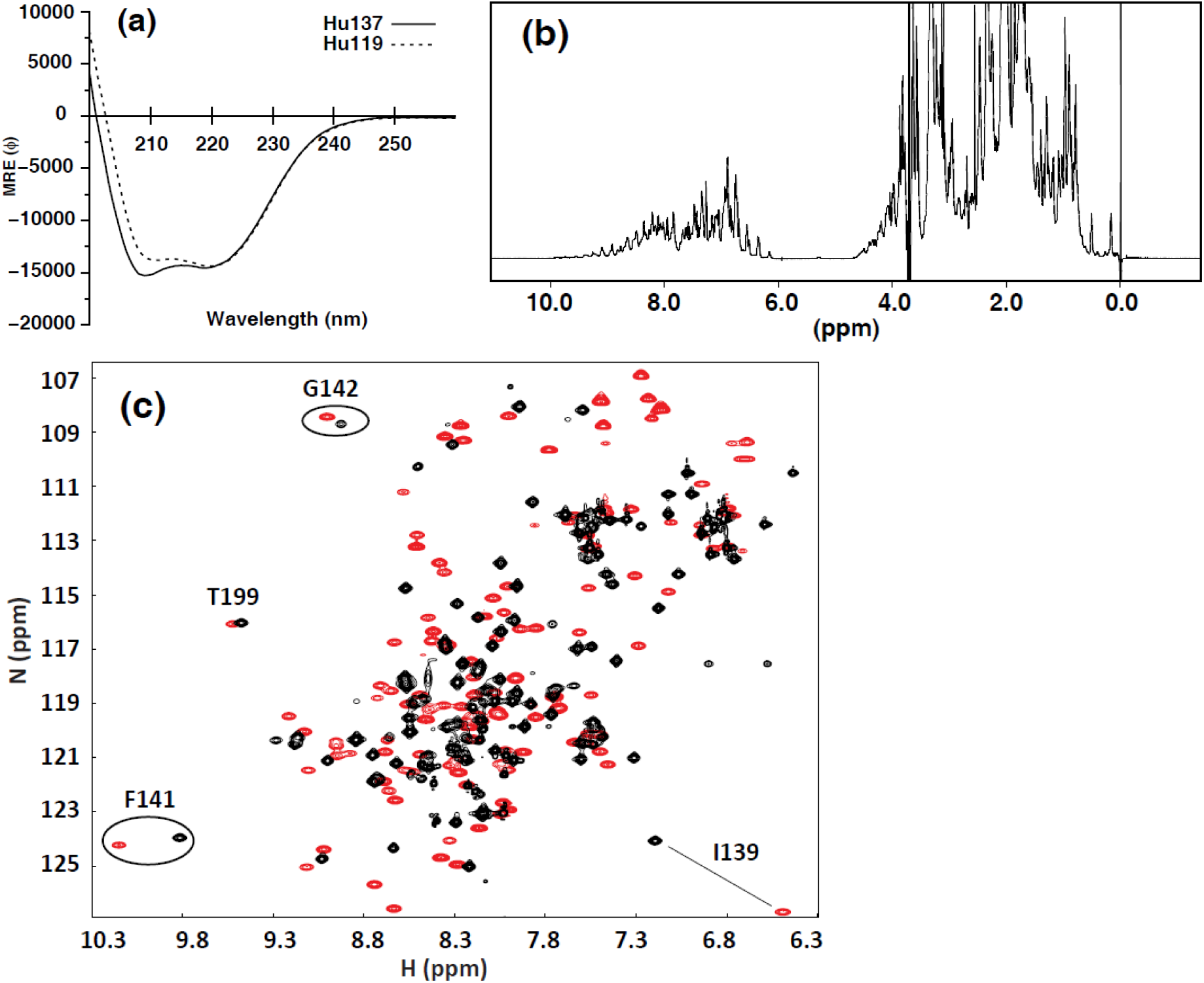
Secondary and tertiary structure of PrP^137^. *(a)* Far UV-CD spectra of PrP^137^ and PrP^119^. Both proteins display CD spectra dominated by α-helical structure, with typical minima at 208 and 222 nm. *(b)* 1D ^1^H NMR spectrum of PrP^137^. The NMR spectrum is typical for a fully folded protein, with well-dispersed NMR signals, including resonances at higher field than 0.7 ppm, characteristic of strong tertiary interactions between methyl groups and aromatic rings, indicating extensive tertiary organisation within the protein. *(c)* Overlay of ^1^H-^15^N HSQC spectra of PrP^137^ (red) and PrP^119^ (black). Signals from regions of the protein in close proximity to the truncation are markedly shifted (I139/F141), in contrast to those remote from the truncation.

Both 1D and 2D NMR spectra of PrP^137^ display a broad range of dispersed NMR signals (Fig. 2 (b/c)). The wide chemical shift dispersion indicates extensive tertiary organisation, and tight packing of amino acid side chains in the protein core characteristic of a fully folded, native-like protein. This is significant, as in so-called ‘molten globule’ intermediate states, truncated or destabilised proteins can display native-like secondary structure content, and compact hydrodynamic radius, despite the absence of well-defined tertiary structure (35). The characteristics found in molten globule states have been found in the transient intermediate states found during the folding of certain proteins (36). However, PrP^137^ NMR spectra are markedly different to PrP^119^, indicating considerable perturbation of the folded domain, in particular residues in close proximity to the N-terminal deletion (Fig. 2 (c)).

### Structural perturbations in PrP^137^ resulting from loss of CHR

The full extent of the structural perturbations resulting from the loss of the CHR was determined through analysis of backbone NMR chemical shifts. Differences between observed chemical shifts and their corresponding random coil (unstructured) values, in particular in Cα and carbonyl resonances, are highly correlated with protein secondary structure (37, 38). Chemical shift data also provides accurate, quantitative and site-specific mapping of protein backbone mobility as well as *Modelfree* backbone order parameters (39, 40).

NMR resonance chemical shifts (Cα/C’/Cβ/N/H^N^) show that the backbone conformation of PrP^137^ is similar to PrP^119^, with all three α-helices retained (Fig. 3) (37, 38). Remarkably, although some loss of structure is observed, the second β-strand also retains clearly observable extended structure despite the loss of the complementary β-strand and its stabilising hydrogen bonding contacts (Fig. 3). A notable change however is an increase of helical structure in the loop linking the second strand of the β-sheet and helix 2, the so-called β2-α2 loop (residues 165–172; Fig. 3). This region, which has been shown to affect prion cross-species transmissibility (41-45), is adjacent to the β-sheet, packing against residues N-terminal to the first β-strand, the β-sheet itself, and the C-terminus of helix 3 (Fig. 1). This structural perturbation appears to have propagated to the C-terminus of helix 3 (residues 222 - 227) which shows some indication of a loss of α-helicity, as does the C-terminus of helix 1 (residues 144 - 154) (Fig. 3).

**Figure 3.**
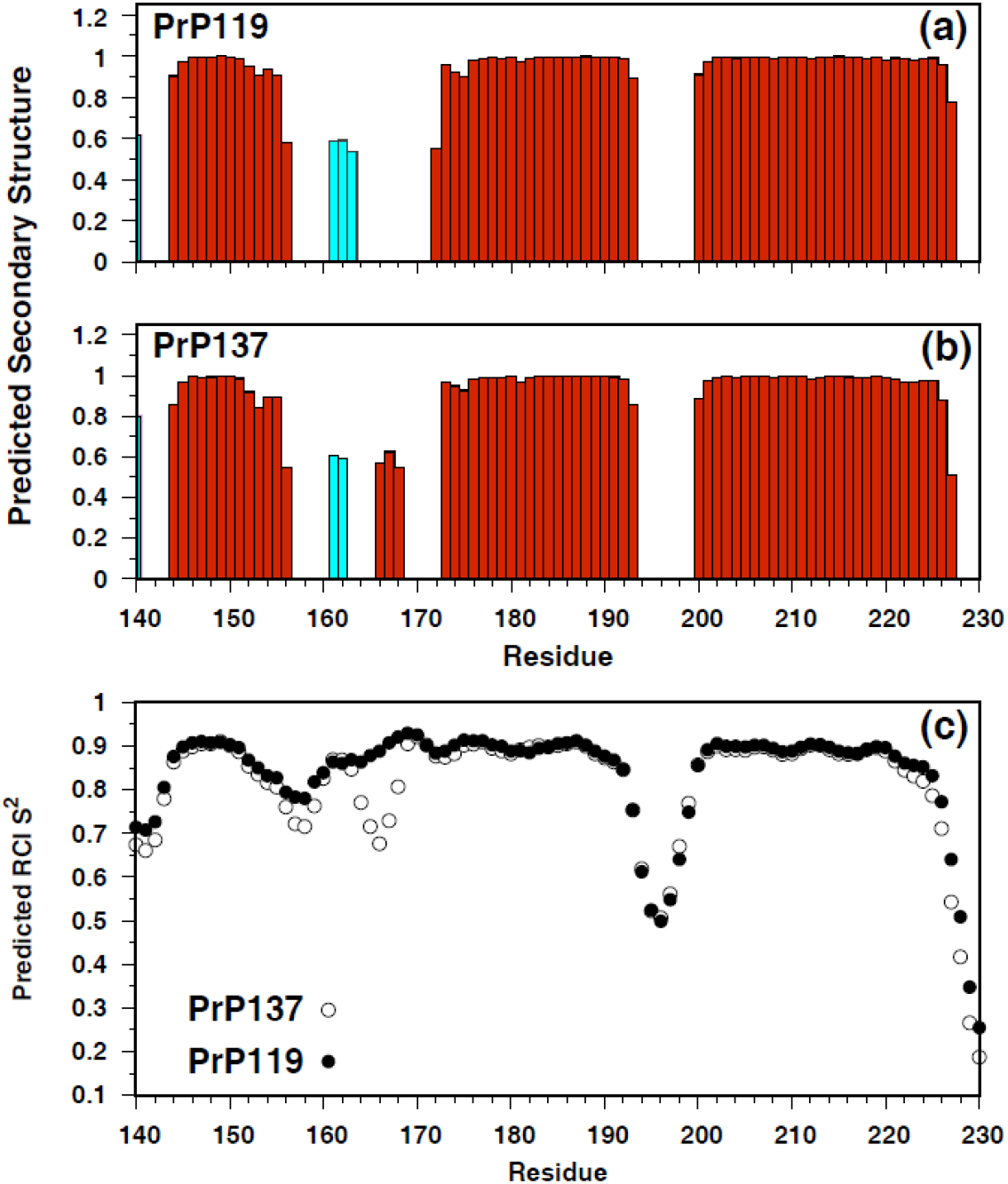
Structural perturbations in PrP^137^ resulting from loss of the CHR. Predicted secondary structure elements for (*a*) PrP^119^ and (*b*) PrP^137^, as calculated by TALOS-N (38). Cyan bars indicate extended β-sheet structure, and red bars α-helical regions. The height of the bars reflects the probability of the structure prediction. (*c*) RCI (Random Coil Index)(40) prediction of order parameter S^2^. S^2^ values report on internal sub-nanosecond (ns) motions, and range from 0 for highly flexible to 1 for rigid systems. An increase in mobility at the C-termini of helices 1 and 3 (residues 144-154; 218-230) and in the β2-α2 loop (residues 166-171) of PrP^137^ is observed.

These alterations in secondary structure propensity are associated with changes in dynamics. For instance the β2-α2 loop displays a marked reduction in the *Modelfree* order parameter (S^2^) (Fig. 3 (c)) (39, 40). S^2^ reports on internal sub-nanosecond (ns) motions, and values range from 0 for highly flexible to 1 for rigid systems. The helical regions of PrP variants exhibit S^2^ values of 0.8 – 0.9, typical of structured regions of folded proteins. Residues of the α2–α3 loop (residues 194–199) and the C-terminus of helix 3 display reduced S^2^ values, reflecting increased flexibility, commonly observed in loop regions of globular proteins (46, 47). The reduction in S^2^ for residues 164-168 in PrP^137^ would suggest that the β2-α2 loop is more flexible in PrP^137^ than PrP^119^. This would appear to be at odds with the predicted increase in α-helicity. However, this loop is subject to millisecond (ms) timescale motions proposed to be associated with a large-scale co-operative conformational change between a 3_10_-helix and a type I β-turn (48, 49). Millisecond timescale motions can result in line-broadening of NMR signals and anomalous order parameter values (46, 47). Loss of the CHR in PrP^137^ does appear to alter the conformational dynamics of the β2-α2 loop and increase the population of the 3_10_ helix. The C-terminus of helix 3 (residue 219 – 230) also displays reduced S^2^ values (Fig. 3 (c)). A number of residues in this region of the protein interact closely with the β2-α2 loop. For example, Y218 and S222 with M166, and I215 and Y218 with Q172. Perturbation of the dynamics of the β2-α2 loop has previously been shown to affect the dynamics of the C-terminus of helix 3 (47), and in PrP^137^ the loss of the CHR and contacts with the β-sheet appear to have propagated beyond the immediate site of the truncation. In addition, the C-terminus of helix 1 and following residues (residues 153-158) also display an increased level of flexibility (Fig. 3 (c)).

### PrP^137^ stability

Despite the relative lack of secondary structure perturbations observed, removal of the CHR and the first β-strand resulted in a marked reduction of approximately 2 kcal mol^-1^ in the thermodynamic stability of PrP^137^ (Fig. 4 (b)). This equates to an approximately 20-fold reduction in the equilibrium between the native and unfolded states (K_(N/U)_ ∼3500) when compared to PrP^119^. As with PrP^119^, loss of secondary structure upon equilibrium denaturation in PrP^137^ occurs in a single co-operative transition, without the formation of any populated intermediate species (Fig. 4 (a)). This unfolding transition is markedly less cooperative than in PrP^119^ (27), but distinct from unfolding transitions demonstrated by molten globule states. The calculated *m-*values, which describe the sensitivity of the folded / unfolded state equilibrium to denaturant, and reflect the increase in solvent exposure of the hydrophobic core as the protein unfolds show that that the degree of hydrophobic exposure in PrP^137^ on unfolding is significantly reduced in comparison to PrP^119^ (Fig. 4 (c)).

**Figure 4.**
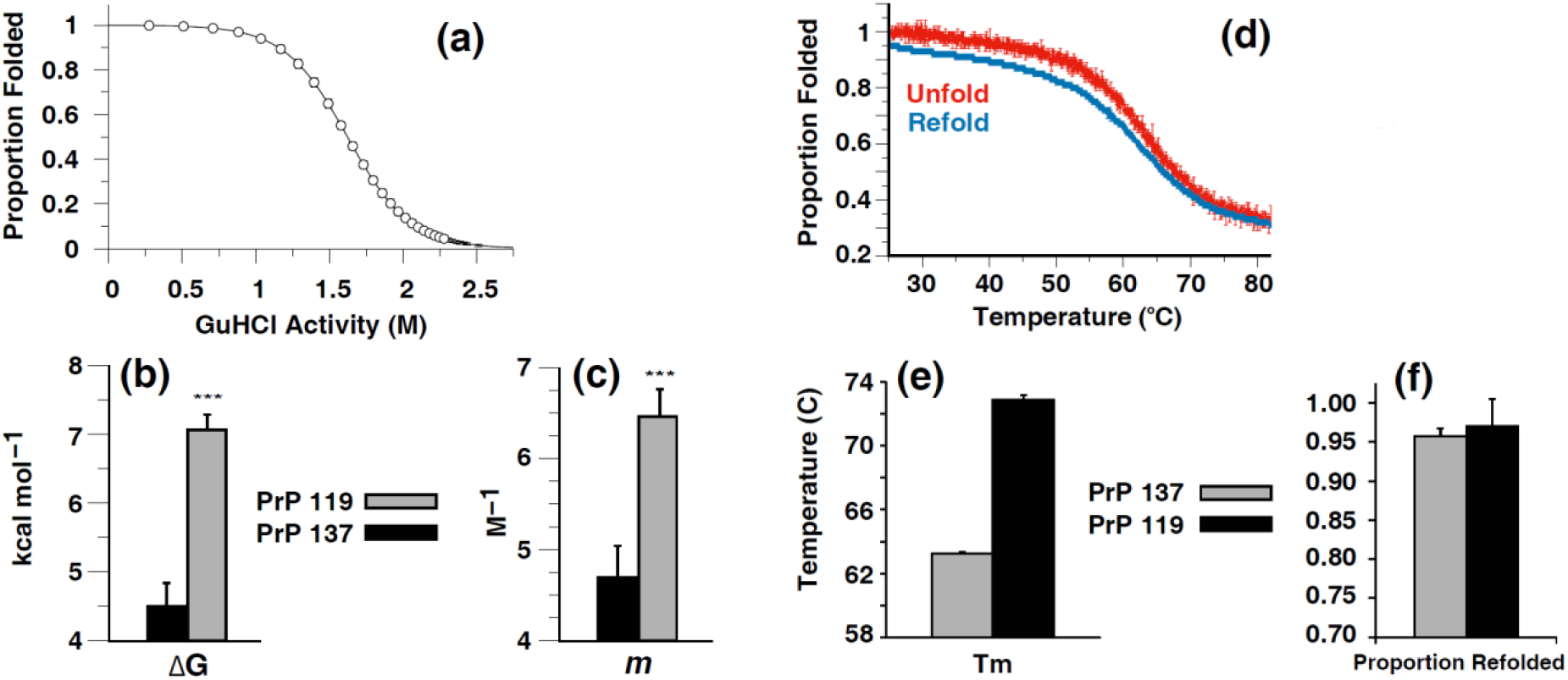
Thermodynamic and thermal stability of PrP^137^. *(a)* Denaturant-induced unfolding transition of PrP^137^. The solid line represents the non-linear least-squares fit to a two-state model of folding (Eqn. 1) with pre- and post-transition baseline slopes. The points shown are corrected for the pre- and post-transition baselines. *(b)* Free energy change (ΔG) of the equilibrium unfolding transition calculated using Eqn. 1. (*c*) Degree of hydrophobic core exposure (*m* values) in PrP^137^ and PrP^119^ derived from the above equilibrium denaturation curve. (*d*) Uncorrected thermally-induced PrP^137^ folding transitions, as calculated by Eqn. 3. *(e)* Temperature mid-point (T_m_) for PrP^137^ and PrP^119^ thermal unfolding transitions. (*f*) Proportion of refolded protein for PrP^137^ and PrP^119^ following thermal denaturation. All folding transitions were monitored by the change in amide CD signal at 222 nm.

In common with the denaturant-induced unfolding, the thermal unfolding of PrP^137^ consisted of a single co-operative transition, without the formation of any populated intermediate species (Fig. 4 (d)). Both equilibrium unfolding transitions (GuHCl- or thermally-induced), were found to be completely reversible (Fig. 4 (d/f)). The thermal stability of PrP^137^ was also markedly reduced in comparison with PrP^119^, with a significant (≈10°C) reduction in the mid-point for thermal unfolding (Fig. 4 (e)).

### Stability of secondary structure elements and solvent accessibility

An even more sensitive measure of local stability is hydrogen/deuterium exchange of backbone amides, observed by the decay of NMR amide signals. As with PrP^119^, protected amides in PrP^137^ were located within the structured elements of the protein, in this case predominantly within helices 2 and 3 (Fig. 5). Protection factors (PF) for amides in these regions were equivalent to the free energy change for unfolding (*i*.*e*. PF = K _(N/U)_). This behaviour is observed in PrP^119^, and also a number of other proteins (*e*.*g*. Barnase/Staphylococcal Nuclease) (50), where a substantial proportion of core residues can exchange only in the fully unfolded state. Protection factors for the second strand of the PrP β-sheet of PrP^137^ were however below detectable levels. This is consistent with the loss of the first strand of the β-sheet and its hydrogen-bonding interactions with the second β-strand.

**Figure 5.**
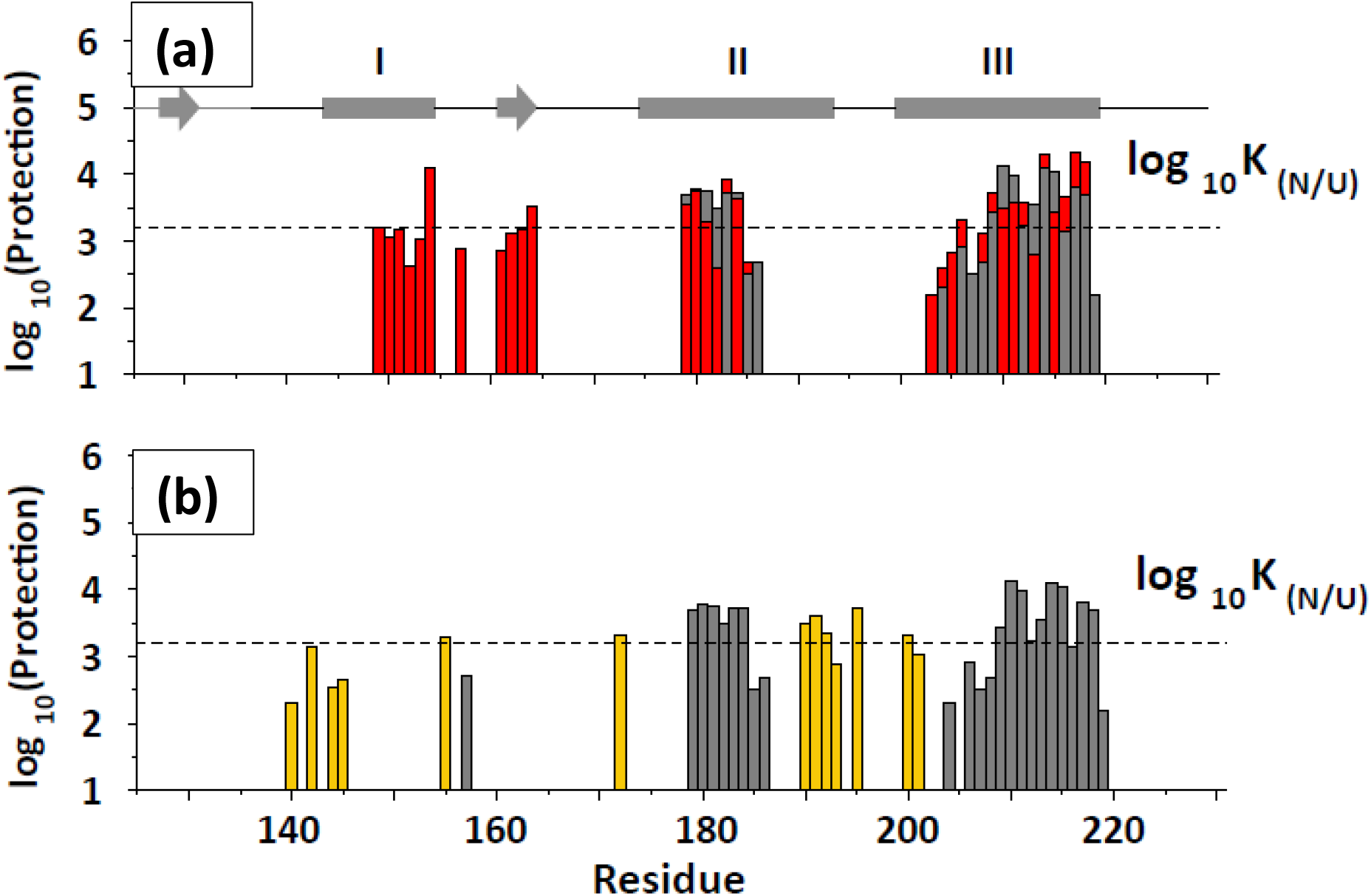
Stability of PrP^137^ secondary structure elements as assessed by native-state hydrogen/deuterium exchange. (a) Amide protection factors (k_ex_/k_int_) in PrP^137^ (grey bars) and PrP^119^ (red bars) for those residues with measurable protection determined through H_2_O/D_2_O exchange. The protection factor corresponding to the equilibrium constant between the native (N) and unfolded (U) states of PrP^137^ is plotted as a dashed line (log_10_ K_(N/U)_). In PrP^119^, the major secondary structure elements (shown at the top of the figure) have stabilities equivalent to the overall stability of the protein, and therefore the protein has to fully unfold for these regions of the protein to exchange with the solvent. In PrP^137^, the second strand of the PrP β-sheet (residues 160-164) does not display measurable protection, consistent with the loss of hydrogen-bonding with the first strand. Notably the first α- helix also displays markedly reduced protection, in comparison with PrP^119^. (b) Amide protection factors for PrP^137^ determined through H_2_O/D_2_O exchange (grey bars/Fig. 5 (a)) and CLEANEX-PM (51) (yellow bars). CLEANEX allows measurement of exchange rates of rapidly exchanging amide protons with lower protection factors, and confirms that that deletion of residues 119 - 136 selectively destabilises helix 1.

However, it was observed that the first α-helix in PrP^137^ displayed markedly less protection than the remaining α-helices. In PrP^137^ it does not display measurable protection (the lower limit for detection of protection factors was approximately 200), whereas in PrP^119^ it has amide protection factors comparable to the other secondary structure elements, and equivalent to the equilibrium constant for unfolding (Fig. 5 (a)). In order to extend the range of measurable protection factors we employed the CLEANEX-PM methodology, which uses phase-modulated chemical exchange NMR to specifically monitor water-amide proton exchange, allowing measurement of exchange rates of rapidly exchanging amide protons (51). This extended the number of measurable amide protons to include residues at the C-termini of helices 2 and 3, and two residues at the N-terminus of helix 1 (Fig. 5 (b)). These displayed protection factors of 350 - 450, approximately 10-fold lower than the average protection factors observed for helices 2 & 3. The remaining residues in helix 1 remained below the detectable limit, indicating that deletion of the CHR has selectively destabilised helix 1.

### Propensity of PrP^137^ to oligomerise

Given the increased exposure of hydrophobic residues, as indicated by the increased *m* value derived from the equilibrium denaturation data and the reduced hydrogen protection for the first α-helix and 2^nd^ β-strand, we sought to determine whether this increased the propensity of PrP^137^ to self-associate. This was initially done using sedimentation velocity analytical ultracentrifugation (SV-AUC) (Fig. 6). A single species with a sedimentation coefficient of 1.6S, characterised by a frictional ratio (*f/f0*) of 1.25, giving a molecular mass of 11.8 kDa was observed for PrP^137^ at pH 7.5. Corresponding values at pH 6.5 were 1.67S, 1.23 and 11.95 kDa respectively. These molecular masses match closely the expected molecular mass for the PrP^137^ monomeric protein (Fig. 6). For PrP^119^ a single species was also observed at pH 6.5 and 7.5 with a molecular mass corresponding with those of the monomeric protein (13.6 kDa). PrP^137^ displays a slightly reduced frictional ratio relative to PrP^119^ indicating a more spherical character, consistent with removal of the relatively unstructured N-terminus (residues 119 - 125) of PrP^119^. Thus it appeared that under the conditions used, PrP^137^ is no more susceptible to oligomerisation than PrP^119^, despite the increased exposure of hydrophobic residues. This conclusion was tested further over a range of protein concentrations (15 – 135 μM) at pH 6.5 and 7.5, with no multimeric species observed (Fig. 6).

**Figure 6.**
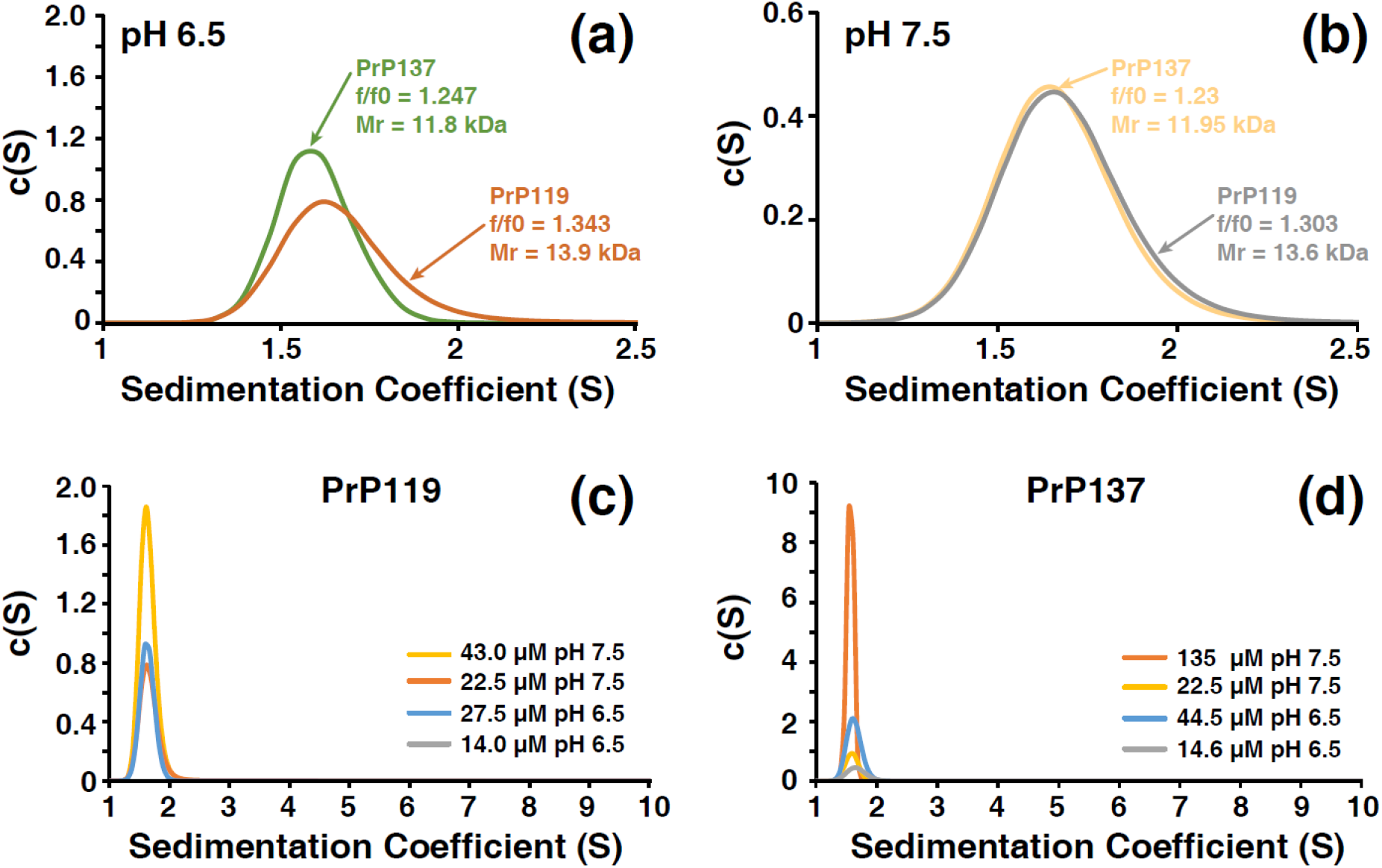
Oligomeric state of PrP^137^ and PrP^119^ determined by sedimentation velocity AUC. Expanded (*a/b*) and full (*c/d*) sedimentation coefficient distributions of PrP^119^ and PrP^137^. (*a/b*) Both PrP constructs sediment with derived molecular masses equivalent to the expected molecular mass of their monomers at both pH 6.5 and pH 7.5. (*c/d*) Protein concentration-dependence for aggregation. Both PrP^119^ and PrP^137^ remain monomeric at pH 6.5 and pH 7.5 over a range of protein concentrations, with no indication of the population of oligomeric states.

### Propensity of PrP^137^ to fibrilise

When agitated at 42°C under native conditions, PrP can be induced to form amyloid (52). Binding of the fluorescent thiazole dye thioflavin T to these β-sheet-rich fibrillar structures reports their formation, allowing a quantitative analysis of the kinetics of fibril formation (53). Although no association of native PrP^137^ monomers was observed in solution, we found that PrP^137^ can be induced to fibrilise much more readily than full-length WT PrP, with significantly shorter half- and lag-times (Fig. 7).

**Figure 7.**
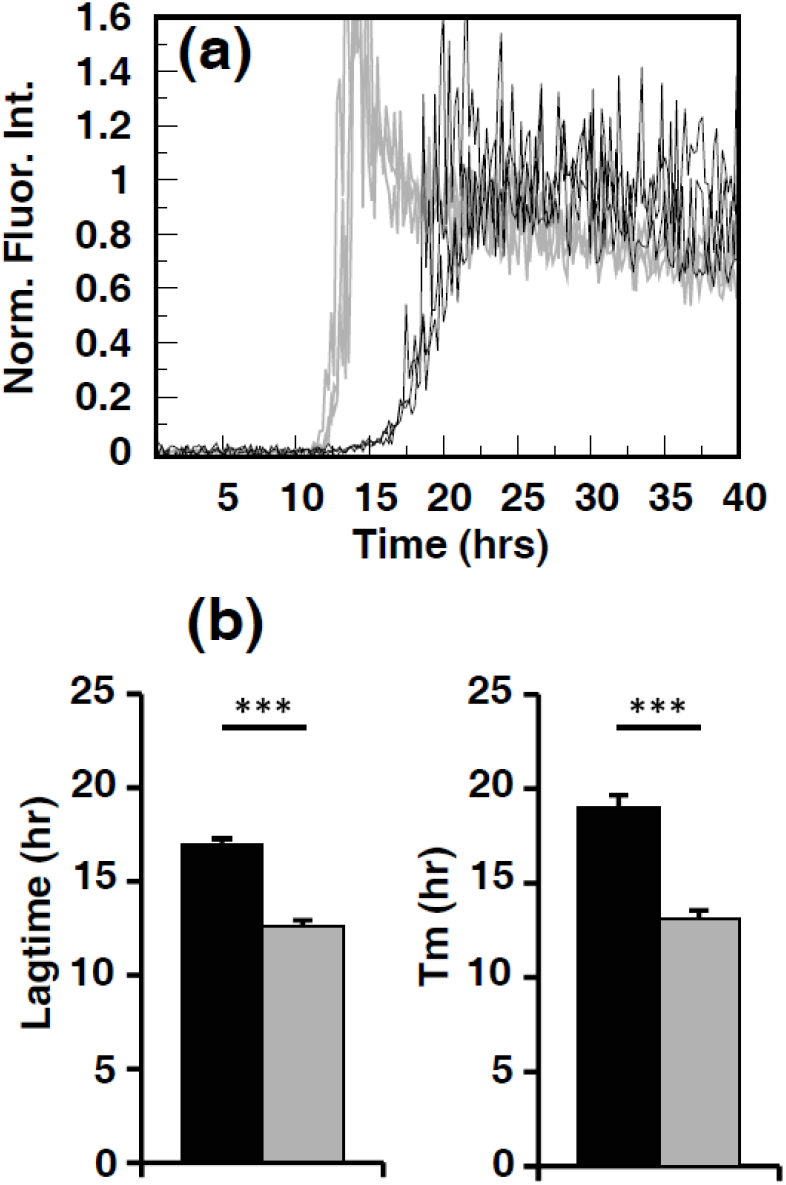
Quantitative analysis of the effect of deletion of the CHR on PrP fibril formation, as reported by increasing Thioflavin T (ThT) fluorescence. (a) Kinetic traces of ThT fluorescence (PrP^137^ – light grey; PrP^23^ – black). Three replicate traces for each protein are shown. (b) Fibrillogenesis of PrP^137^ occurs with significantly longer mean half- and lagtimes in comparison to PrP^23^ (P ≤ 0.001, paired t-test). The data shown are the mean readings from five replicates ±SD.

## Discussion

In this study, we have examined the effect of the removal of the N-terminus of the folded domain of PrP, on the structure, stability and fibrillisation characteristics of PrPC. This truncated form of PrPC mimics the structural effect of the association of the conserved hydrophobic region with the ER membrane on the folded C-terminal domain of PrP^C^.

We find that the truncated protein retains the native secondary structure elements of PrP^C^, including, remarkably, the second strand of the PrP β-sheet, but with increased conformational flexibility at the C-termini of α-helices 1 and 3 and altered dynamics in the conformationally-variable β2-α2 loop. It also retains a well-defined tertiary structure, with strong evidence for co-operative folding. There is a marked reduction in the stability and increased exposure of hydrophobic residues however, which would appear to facilitate its conversion to fibrillar states significantly. The structural elements of PrP^C^ most affected by the deletion are the second strand of the β-sheet and the first α-helix, both of which display markedly increased solvent exposure and reduced stability relative to the other structure elements.

Previous hydrogen exchange data show that the core of PrP^C^, comprising the three α-helices and the second β-strand, is a stable entity, which has to completely unfold for exchange with the solvent to occur (26). These data suggest that for fully folded PrP^C^, the most likely route to PrP^Sc^ is through the completely unfolded state, and that any partially-folded intermediate states involved in this conversion would most likely retain this core region.

Under partially denaturing, acidic conditions however, various regions of mouse PrP^C^ have been shown to undergo sub-global unfolding, forming at least two distinct partially unfolded forms (PUFs) (54). These are in equilibrium with the native state and display increased solvent exposure relative to it. A key characteristic of the second of these PUFs is that the first α-helix, second strand of the β-sheet and loop regions in between are disordered and solvent accessible. In this study we find that that removal of the PrP N-terminus to residue 137, mimicking the insertion of the CHR into the ER membrane, also markedly reduces the stability of α-helix 1 and the second β-strand, generating a partially unfolded form of PrP^C^ which displays many of the characteristics of the PUF generated under partially-denaturing acidic conditions (54). Both of these open forms fibrillise much more readily than the native form of the protein, and resemble a conformation implicated to be an initial intermediate in the conversion of monomeric PrP into misfolded oligomer at pH 4 (55). α-helix 1 in particular has been implicated in the conversion to PrP^Sc^ (56, 57), with the anti-PrP therapeutic monoclonal antibody ICSM18 effectively curing prion-infected cells through binding to and stabilising α-helix 1 of PrP^C^ (58). Indeed its humanised version has been used in the first-in-human treatment programme using an anti-PrP^C^ monoclonal antibody (59).

The majority of the CHR itself is largely unprotected from amide exchange in PrP^C^ (26), including the first PrP β-strand, which displays anomalously low protection factors in PrP^C^. This is intriguing given the full protection seen in the other paired β-strand. The detachment of the CHR from the PrP core thus raises two possibilities, firstly that the PrP β-sheet is maintained under native conditions and the loop regions not including the first β-strand (residues 125 – 127, 132 - 145) detaches, or that secondly, the residues within the β-sheet that are involved in hydrogen bonding within the PrP β-sheet are involved in hydrogen bonding with other hydrogen bond acceptors. The marked reduction in protection for the second β-strand in PrP^137^ would suggest that the first possibility is more likely, at least in solution.

Overall, our results are concordant with other *in vitro* experiments that show that destabilisation of PrP^C^ favours the formation of fibrillar and/or protease resistant structures (60, 61), a number of which have been proposed to be associated with the disease process (23-25). PrP^C^/PrP^Sc^ conversion may thus be initiated or facilitated by an increased population of this aggregation competent state, formed by insertion of the CHR into the ER membrane. This proposal is consistent with the observation that antibody sequestration of the CHR inhibits mouse prion propagation (34). This may be through sequestration of the CHR from membrane insertion and or stabilisation of the PrP^C^ native state.

Our data suggest an alternative pathway to structural conversion of PrP to the prion conformer, in which the detached CHR residues destabilize the folded core of the protein, thus lowering the energy barrier to its structural conversion. The CHR could interact with the active surface of the prion fibril first, thus destabilizing the rest of the protein core and generating the “open” form characterised here, which refolds afterwards. Alternatively, destabilization by the membrane-bound CHR could facilitate interaction of α-helix 1 with the prion, initiating the structural conversion process. The data presented thus highlight the key role of the central hydrophobic region of the prion protein on the folded domain of PrP^C^, and provide an intriguing explanation to how this may facilitate the conversion of PrP^C^ to PrP^Sc^.

## Materials and Methods

### Protein Expression and Purification

The open reading frame of the human PrP gene (PRNP) (residues 137-231), containing methionine at residue 129, was synthesised *de novo* by *Eurofins MWG Operon*, with a thrombin-cleavable His-Tag added to the PrP N-terminus. The ligated pTrcHisB/PRNP construct was used to transform the *Escherichia coli* host strain BL21(DE3) (*Novagen*), genotype *F′ ompT hsdSB (rB-mB-) gal dcm* (DE3), which was then plated onto Luria-Bertoni (LB) agar plates containing 100 μg/ml carbenicillin. Cultures were grown for purification using a modification of protocols previously described (27). Briefly, following harvesting, cells were sonicated and their inclusion bodies containing PrP resolubilised in 6M Guanidine Hydrochloride (GuHCl), 50 mM Tris.Cl, 0.8% β-mercaptoethanol, pH 8.0. These were loaded onto a Ni-NTA column equilibrated in 6 M GuHCl, 10 mM Tris-HCl, 100 mM Na_2_PO_4_, 10mM glutathione pH 8.0, and eluted from the column using 10mM Tris-HCl, 100 mM Na_2_PO_4_, 1M Imidazole pH 7.0. Residual GuHCl was removed through dialysis against 25 mM Tris.HCl pH 8.4, CaCl_2_ added to a final concentration of 2.5 mM, and the N-terminal His-tag cleaved by thrombin for 16 h at room temperature (0.1 U thrombin (*Novagene*)/1 mg of PrP added). The cleaved protein was loaded onto Q-Sepharose column equilibrated with 10 mM HEPES pH 8.2, and the PrP eluted by the same buffer containing 1M NaCl. PrP^137^ was dialysed against 10 mM HEPES pH 7.5, and stored at -80°C. Protein concentrations were determined by UV absorption using a calculated molar extinction of 15025 M^-1^ cm^-1^ at 280 nm (*https://web.expasy.org/protparam*).

### Circular Dichroism (CD) spectroscopy

CD spectra were measured with a Jasco J-715 spectropolarimeter. Far-UV (amide) CD spectra (300 – 180 nm) were acquired on 22.5 μM protein at 25°C using 0.5 mm pathlength quartz cuvettes. The sample temperature was controlled with a circulating water bath. 45 spectra were averaged.

### Equilibrium unfolding experiments

6.5 µM PrP^137^ and PrP^119^ in 10 mM sodium phosphate, 1 mM sodium azide, pH 7.0, were incubated in increasing concentrations of GuHCl denaturant at 25°C. Molecular ellipticity ([θ], degree M^-1^ cm^-1^) was recorded at 222 nm (5 nm bandwidth; 20 s integration time) in the Jasco J-715 spectropolarimeter. The denaturation profile for each protein was measured in 3 separate experiments.

### Calculating the equilibrium constant between folded and unfolded states (K & K_w_)

For the two-state equilibrium unfolding transitions, data were fitted to the following equation, where K and K_(W)_ are equilibrium constants between the folded and unfolded states at a given denaturant activity (D) and in water, respectively, and *m* describes the sensitivity of the equilibrium to denaturant activity (62).

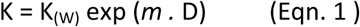

For visual representation of the data shown, data were converted to proportion folded, α_F_, using the following, α_F_ = (K/(1+K)). Data fitting was carried out using *GNUplot* v. 5.2. The significance of the differences in free energy for folding and *m* values between PrP^119^ and PrP^137^ were determined by two-tailed student’s t-Test.

### Calculation of Denaturant Activity

Due to the non-linear relationship between denaturant concentration and the free energy for folding, GuHCl concentration ([GuHCl]) was converted to molar denaturant activity to obtain a more reliable extrapolation of data at high denaturant concentrations. The equation used is

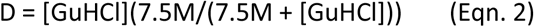

where D is the molar denaturant activity (62).

### Equilibrium Thermal Denaturation Monitored by CD

The amide CD absorption of 6.5 μM PrP^137^ in 10 mM HEPES, 25 mM NaCl, 1 mM sodium azide, pH 7.0, was recorded at increasing temperature (1 °C/minute change). The ellipticity signal (α) was converted to the proportion of molecules in the native state α_N_ according to the relationship α_N_ = (θ - θ_U_)/(θ_N_ - θ_U_), where θ_U_ and θ_N_ are the ellipticity signals for the unfolded and native states, respectively. The Van’t Hoff enthalpy (H), temperature mid-point for thermal folding (T_m_), and equilibrium constant for folding (K_(FU)_)were calculated as follows.

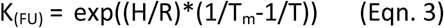

where R is the gas constant (kcalmol^-1^K^-1^). All equilibrium unfolding transitions, GuHCl- or thermally induced, were reversible.

### NMR spectroscopy

Uniformly ^15^N- and ^13^C/^15^N-labeled PrP^137–231^ samples were expressed in *E. coli* using an EMBL minimal medium recipe with ^13^C_6_-glucose and (^15^NH_4_)_2_SO_4_ as the sole carbon and nitrogen sources, respectively, and purified as described above. NMR spectra were acquired at 293K on 1.0 mM ^15^N- and ^13^C/^15^N-labeled samples in 10 mM sodium phosphate, 2 mM sodium azide, 1mM TSP, pH 7.0 (in either 90% H_2_O plus 10% D_2_O *(v/v)* or 50% D_2_O *(v/v)*) using Bruker DRX-500 and DRX-600 spectrometers equipped with 5-mm ^13^C/^15^N/^1^H triple resonance probes. NMR samples were placed in Sigma FEP NMR sample tube liners (Z286397-1EA) held within Wilmad PP-528 NMR tubes for NMR data acquisition. Proton chemical shifts were referenced to TSP, with ^15^N and ^13^C chemical shifts calculated relative to TSP, using the gyromagnetic ratios of ^15^N, ^13^C, and ^1^H (^15^N/^1^H = 0.101329118, ^13^C/^1^H = 0.251449530). NMR data were processed and analysed on Linux Workstations using *Felix 2007* (Accelrys, San Diego) software.

### NMR backbone assignments

#### Backbone NMR assignment

Backbone resonances (^H^N, N, Cα, C′, Cβ) of PrP^137^ were obtained using a standard suite of triple resonance NMR experiments (63), and assigned using the *asstools* set of assignment programs (64). Almost complete backbone assignments were determined, the exceptions being the amide protons and nitrogens of residues 167, 169–171, and 175, for which no resonances were detected. These residues occupy a loop region between β-strand 2 and α-helix 2, which is undergoing conformational exchange, resulting in line-broadening of NMR signals (47).

#### Prediction of secondary structure propensity and ModelFree Analysis

Secondary structure propensity was calculated from H^N^, Cα, Cβ, CO, & N random coil chemical shifts and average secondary shifts for protein secondary structure using the program TALOS-N (38). TALOS-N is an artificial neural network based system for empirical prediction of protein backbone φ/ψ torsion angles, sidechain χ1 torsion angles and secondary structure using chemical shifts. The *Modelfree* order parameters S^2^ for the backbone amide groups were predicted from backbone (H^N^, Cα, CO, & N) and Cβ chemical shifts using the Random Coil Index (RCI) approach as implemented within TALOS-N (38-40).

#### Amide Hydrogen-Deuterium Exchange

Hydrogen-deuterium exchange rates (k_ex_) were determined by adding 140 µl 10 mM sodium phosphate, 1 mM sodium azide, pH 7.0, dissolved in 100% (*v/v*) D_2_O to the same volume of 1 mM PrP^137^ in the equivalent protonated buffer. A series of 2D sensitivity-enhanced ^1^H-^15^N HSQC spectra (65, 66) were acquired at 293K on a Bruker DRX-800 spectrometer. The decay curves of the ^1^H-^15^N HSQC cross-peaks were fitted to single exponential decays with offset, and protection factors (k_ex_/k_int_) for observable amides were determined using intrinsic amide exchange rates (k_int_) (67). Acquisition of the first experiment began approximately 5 minutes after mixing.

#### Amide Hydrogen-Deuterium Exchange measured by CLEANEX

Fast exchanging amide proton rates were determined using the CLEAN chemical EXchange (CLEANEX-PM) approach (51) under the same conditions used for the hydrogen-deuterium exchange experiment described above.

#### Analytical Ultracentrifugation

Sedimentation velocity ultracentrifugation experiments (SV-AUC) were carried out using a Beckman Optima XL-I analytical ultracentrifuge. Samples were loaded into Beckman AUC sample cells with 12 mm optical path two-channel centrepieces, with matched buffer in the reference sector. Cells were spun at 50,000 rpm in an AnTi-50 rotor and scans were acquired using both interference and absorbance optics (at 280 nm) at 10 minute intervals over 16 hours. The sedimentation profiles were analysed using the software SEDFIT (v13b) (68). Partial specific volumes (ρ) for PrP^137^ were calculated from the amino acid sequence using SEDNTERP software (69). Buffer densities and viscosities were measured using an Anton Paar DMA5000 density meter and an Anton Paar AMVn automated microviscometer, respectively. Sedimentation velocity data were analysed using the *c(s)* method of distribution (68) to characterise the sedimentation coefficient distribution of all species present in solution. The proportions of each sample occupying the main peaks in the distribution were calculated by integration of the peaks.

#### Quantitative analysis of the kinetics of PrP fibril formation

PrP^137^ and PrP^23^ were diluted to 5μM in 125mM Sodium Phosphate, 150mM NaCl, 7.5mM Bis-Tris pH 6.7. All solutions were filtered through a 0.22 µm filter to remove particulates. 100 µl aliquots were placed in Greiner 96-well flat-bottomed plates (#655077) containing 3 0.5 mm diameter zirconium ceramic beads in each well to assist agitation. The plates were incubated at 42 °C with constant agitation in a Tecan Infinite F200 Microplate Fluorimeter. Fibril formation was monitored through the increase in ThT fluorescence (excitation 430 nm, emission 485 nm, 20 nm bandwidths), with readings acquired every 600 seconds. 3 replicates were used for each PrP sample.

To determine the half- and lag-times for fibril formation, data were fitted to an empirical function described by Nielsen et al (2001)(70).

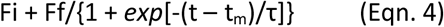

where Fi is the initial fluorescence reading, Ff is the final fluorescence reading, t is time, t_m_ is the time taken to half maximal fluorescence and τ is the reciprocal of the propagation rate during the rise phase [1/k_(apparent)_]. Lag-time is defined as t_m_ - 2τ.

## Abbreviations

BSE: bovine spongiform encephalopathy
CD: Circular Dichroism
CJD: Creutzfeldt-Jakob disease
FFI: fatal familial insomnia
GuHCl: guanidine hydrochloride
MRE: Mean Residue Ellipticity
PrP: prion protein
PrP^C^: cellular PrP isoform
PrP^Sc^: pathogenic (scrapie) PrP isoform
HuPrP: human PrP
PrP^23/119/137^: residues 23/119/137–231 of human PrP
HSQC: heteronuclear single quantum coherence
SV-AUC: sedimentation velocity analytical ultracentrifugation
TSP: sodium 3-trimethylsilyl-2,2,3,3-(^2^H_4_) propionate.

